# Pathogenesis of *Achromobacter xylosoxidans* respiratory infections: colonization and persistence of airway epithelia and differential gene expression in synthetic cystic fibrosis sputum medium

**DOI:** 10.1101/2023.04.04.535650

**Authors:** Caitlin E. Billiot, Melissa S. McDaniel, W. Edward Swords, Natalie R. Lindgren

**Author notes:** Communicating author: 1918 University Boulevard, MCLM 818 Birmingham, AL 35294. Department of Microbiology University of Alabama at Birmingham (Melissa S. McDaniel).

## Abstract

Cystic fibrosis (CF) is a genetic disease affecting epithelial ion transport, resulting in thickened mucus and impaired mucociliary clearance. Persons with CF (pwCF) experience life- long respiratory mucosal infections caused by a diverse array of opportunists, and these infections are a leading cause of morbidity and mortality for pwCF. In recent years, there has been increased appreciation for the range and diversity of microbes in CF-related respiratory infections. Introduction of new therapeutics and improved detection methodology has revealed CF related opportunists such as *Achromobacter xylosoxidans* (*Ax*). *Ax* is a Gram-negative bacterial species that is widely distributed in the environment and has been increasingly observed in sputa and other samples from pwCF; typically *Ax* infections occur in patients in later stages of CF disease. In this study, we characterized CF clinical isolates of *Ax* and tested colonization and persistence of *Ax* in respiratory infection using immortalized human CF respiratory epithelial cells and BALB/c mice. Genomic analyses of clinical *Ax* isolates showed homologs for factors involved in flagellar synthesis, antibiotic resistance, and toxin secretion systems. *Ax* isolates adhered to polarized CFBE14o- human immortalized CF bronchial epithelial cells and caused significant cytotoxicity and depolarization. *Ax* colonized and persisted in mouse lung for up to 72 hours post infection, with inflammatory consequences that include increased neutrophilia, lung damage, cytokine production, and mortality. Transcript profiling reveled differential expression of *Ax* genes during growth in SCFM2 synthetic CF sputum media. Based on these results, we conclude that *Ax* is an opportunistic pathogen of significance in CF.

## INTRODUCTION

Cystic fibrosis (CF) is a genetic disease caused by mutations affecting the function of an epithelial ion channel, the cystic fibrosis transmembrane conductance regulator (CFTR) (1–3). Of note, persons with CF (pwCF) have thickened mucus and decreased mucociliary clearance, which leads to chronic lung infections that remain a leading cause of morbidity and mortality in pwCF. While treatment options for CF have improved in recent years, there is no standard antibiotic treatment for CF related lung infections (4, 5). With the introduction and widespread use of 16S rRNA sequencing, there is an increased appreciation for the diversity of the CF microbiome, leading to the identification of organisms undetectable by traditional culture-based methods (6, 7). This, with recent increases in the median age for the CF patient population and widespread use of broad-spectrum antibiotics, there has been an increase in non-traditional CF pathogens that are late-stage colonizers, such as *Achromobacter xylosoxidans* (*Ax*) (8, 9). *Ax* is a Gram-negative, motile, and inherently antibiotic resistant bacterium that causes opportunistic infections such as pneumonia, meningitis, and urinary tract infections (9). *Ax* has been recognized as a CF related opportunist since the early 1980’s, and since then, has been regarded as an emerging CF pathogen, as it is increasingly isolated from pwCF worldwide (9, 10).

Despite increasing prevalence, little is known about how this organism affects the disease progression in pwCF. Genomic sequencing of *Ax* isolates has revealed genes associated with pathogenesis in other bacterial species, but virulence factors encoded by these genes have not been examined experimentally in *Ax* (9). These virulence factors include: the Vi capsular polysaccharide, type III secretion systems, multi-drug efflux pumps, and denitrification and iron chelation systems, all of which have been shown to provide a survival advantage during host infection in other bacterial species (9, 11). Despite indication that *Ax* carries a wide arsenal of virulence factors, this pathogen is largely understudied *in vivo* and its effect on CF disease progression is not well understood.

Epidemiological studies using CF clinical isolates showed that infection with *Ax* may have clinical significance. For example, it has been demonstrated that patients infected with *Ax* have a lower forced expiratory volume in 1 second (FEV_1_), at baseline when compared to non-colonized patients, with a larger decline in FEV_1_, and higher pulmonary exacerbation frequency for up to 3 years after the first incidence of *Ax* isolation (12). Additionally, another study has demonstrated that chronic *Ax* infection is associated with greater risk of death or lung transplantation compared to those with no history of *Ax* infection, and that chronically infected pwCF had a lower FEV_1_ (13). However, there are also indications that respiratory symptoms in these chronically infected individuals did not significantly worsen after the development of chronic *Ax* infection (13).

These studies provide some insight on the impact of *Ax* on the health of pwCF, but our understanding of specific virulence factors and how they play a role in the pathogenesis of this organism *in vivo* is limited. Recent studies have identified *Ax* genes associated with biofilm formation (14), and macrophage cytotoxicity (15, 16). In this study we aimed to examine pathogenic phenotypes in *Ax* as well as establish a pulmonary infection model to further study the consequences of *Ax* infection. We also aimed to identify specific genes or gene sets that may be important for *Ax* pathogenesis or survival in the CF lung environment through transcriptome profiling of *Ax* grown in a synthetic CF sputum media that is made to mimic the chemical composition of the CF lung.

## RESULTS

### Comparative analyses of cystic fibrosis isolates of *A. xylosoxidans*

Previous literature shows that *Ax* strains can express a multitude of virulence factors, including but not limited to flagellar synthesis genes, biofilm components, pili and fimbriae related genes, and toxin secretion systems. We performed preliminary genomic analysis to compare our clinical isolates and saw that each of our isolates contain homologs to previously identified virulence genes (data not shown). To examine differences between *Ax* clinical isolates, we performed *in vitro* assays to examine some hits from genotypic analyses, as well as to test pathogenic phenotypes seen in previous literature using three *Ax* clinical isolates from CF patient sputum. We first examined biofilm formation on an abiotic surface and different types of motility (swimming, swarming, and twitching). *Ax* 1 and *Ax* 7 form statistically different amounts of biofilm matrix at all timepoints except for the 24-hour timepoint (Fig. S1A). Bacterial enumeration of these strains at each timepoint, however, are not significantly different between strains at each timepoint (data not shown). Because *Ax* is known to be motile, a concept supported by the annotation of flagellar and pili/fimbriae genes in our genomic analyses, we examined motility. Despite all having the presence of flagellum related genes, *Ax* 1 and *Ax* 7 did not exhibit swimming motility at either of the time points, while *Ax* 10 swam significantly more than those two strains (****P<0.0001) (Fig. S1B). We examined swarming motility of each isolate because this type of motility is also mediated by the presence of flagellum. Notably, none of the isolates examined here exhibited swarming motility (data not shown). Twitching motility is primarily mediated by fimbriae or pili and was also assessed. We observed that each of the strains examined exhibits twitching motility, and that the extent of this motility does not significantly differ between strains at each time point (Fig. S1C).

### *A. xylosoxidans* isolates differentially adhere to immortalized human CF bronchial epithelial cells

We next examined *Ax* adherence to human respiratory epithelia. Using a cystic fibrosis bronchial epithelial cell line homozygous for the F508del mutation (CFBE41o-, hereby referred to as CFBEs), we performed adherence assays on submerged cell cultures using a multiplicity of infection (MOI) of 10 *Ax* cells per mammalian cell at a one hour timepoint, and expressed as percent of inoculum adhered (Fig. 1A). We observed that *Ax* 1 and *Ax* 7 adhere significantly more (** P<0.01) to submerged CFBEs than *Ax* 10 (Fig. 1A). Additionally, we examined growth of *Ax* strains on submerged CFBEs at longer timepoints. We inoculated cells with a MOI of 10 of each *Ax* strain, and incubated infected cells for 6 hours, plating for viable colony counting at 2, 4, and 6 hours post infection. Like the simple adherence experiments, we observed less *Ax* 10 CFUs (compared to those of *Ax* 1 and *Ax* 7) at each timepoint post infection (Fig. 1B).

**Figure 1:**
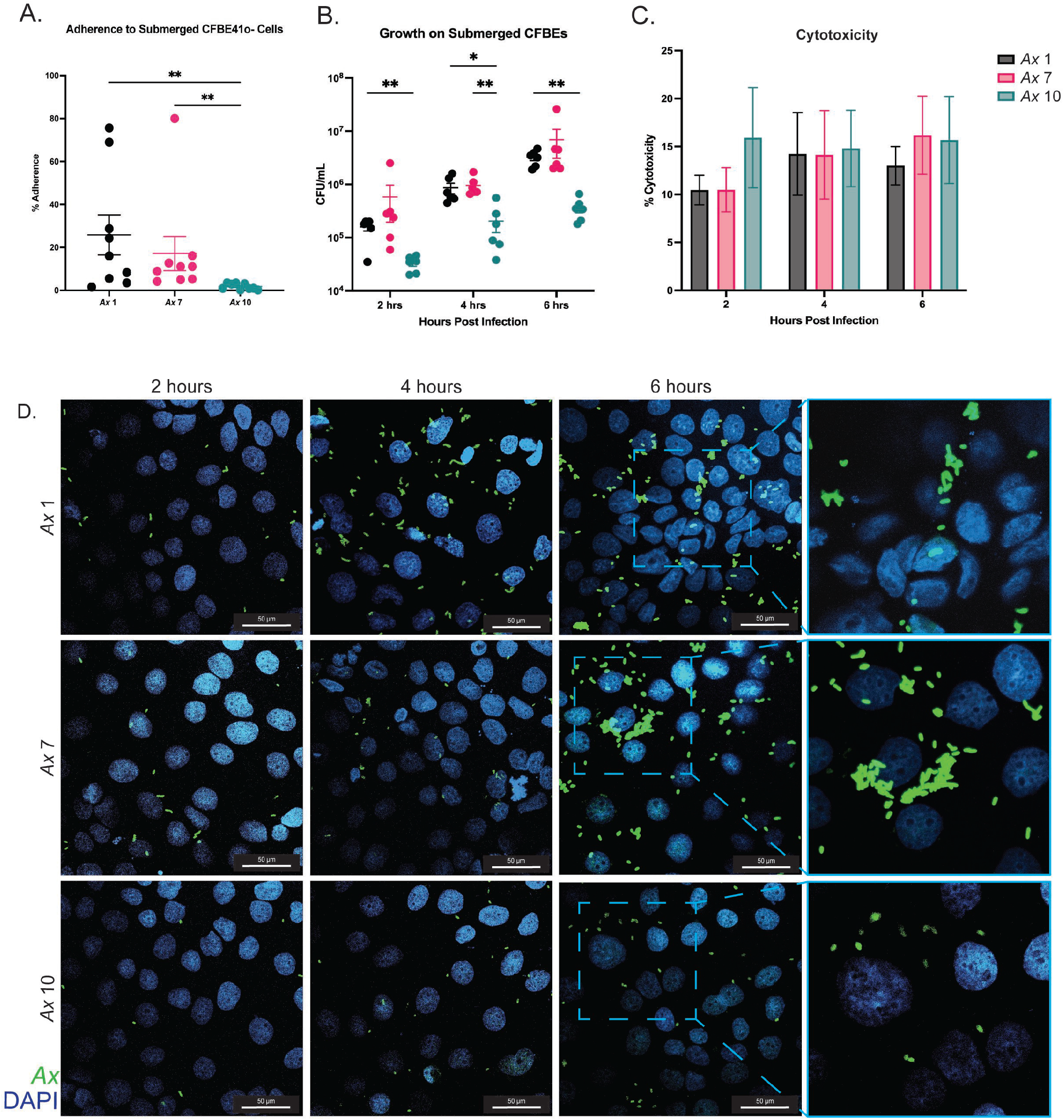
*Ax* strains adhere to cystic fibrosis bronchial epithelial cells and exhibit cytotoxicity. *In vitro* cell culture experiments were completed to investigate *Ax* cytotoxicity and adherence when infecting submerged CFBE41o- (CFBE) cells. CFBEs were infected with MOI 10 of *Ax* and various endpoints were examined. A) Adherence to submerged CFBE cells was examined at the one hour timepoint as expressed by percentage of inoculum. Mean ± SEM. Kruskal-Wallis test with Dunn’s post hoc comparisons. B) Growth of *Ax* strains on CFBE cells was examined at 2, 4, and 6 hours post infection. C) Cytotoxic effects of *Ax* strains on submerged cell culture was examined at 6 hours post infection. Mean ± SEM. Kruskal-Wallis test with Dunn’s post hoc comparisons. * P<0.05, ** P<0.01, *** P<0.001, **** P<0.0001. D) Representative images of CFBEs infected with each strain at each timepoint. Magnification=60x; outlined sections were enlarged with 2x zoom.

To test for cytotoxic effects, we compared CFBE viability at varying time points post infection. Submerged CFBE cells were inoculated with an MOI of 10 of each bacterial strain for 6 hours, and cell supernatant was collected every two hours to perform a lactate dehydrogenase assay (LDH), which was used to calculate percent toxicity. We observed cytotoxicity at each time point for each strain as compared to uninfected controls, and cytotoxicity was not significantly different between strains or time points (Fig. 1C). Additionally, we performed microscopy on infected submerged cells at each time point mentioned previously. CFBEs grown in chamber slides were infected with an MOI of 10 of each *Ax* strain and cells were washed to get rid of non-adherent bacteria and fixed for IF staining. At each time point post infection, *Ax* cells were present on the CFBEs (Fig. 1D). Like the adherence and growth experiments on submerged cells (Fig. 1A and 1B), greater quantities of *Ax* 1 and *Ax* 7 were observed via confocal microscopy at each timepoint post infection. Furthermore, at the 6-hour timepoint, *Ax* 1 and *Ax* 7 form bacterial aggregates, indicating that biofilm formation may be beginning to take place for these strains (Fig. 1D).

### *A. xylosoxidans* infection disrupts epithelial tight junctions in polarized CF epithelia

After observing adherence to CFBE cells grown in submerged culture and growth on these cells, we examined impacts of *Ax* infection on CFBE cells grown at air-liquid interface (ALI) to promote establishment of polarized cell layers and epithelial differentiation (17). Like the submerged cell culture experiments, CFBEs grown at ALI were inoculated with MOI 10 of *Ax* strains. To examine tight junction integrity, measurements of transepithelial electrical resistance (TEER) were taken, and cells were incubated with *Ax* isolates for up to six hours post infection, with TEER being measured every 2 hours. We observed that *Ax* 1 significantly lowered TEER measurements at each time point post infection (Fig. 2B). Additionally, *Ax* 7 significantly lowered TEER measurements at the 2- and 4-hour timepoints, while *Ax* 10’s effect on TEER was non-significant at each of the time points (Fig. 2B). To visualize the tight junction integrity of infected CFBEs grown at ALI, we immunofluorescently (IF)-stained 6-hour infected ALIs for *Ax* and ZO- 1, a component of tight junctions (Fig. 2A). We observed that for each strain, tight junctions appeared to be less intact than that of an uninfected control well (Fig. 2A). We also observed bacterial aggregates in these infections, which appear to be localized to areas in which integrity of the tight junctions was compromised (Fig. 2A).

**Figure 2:**
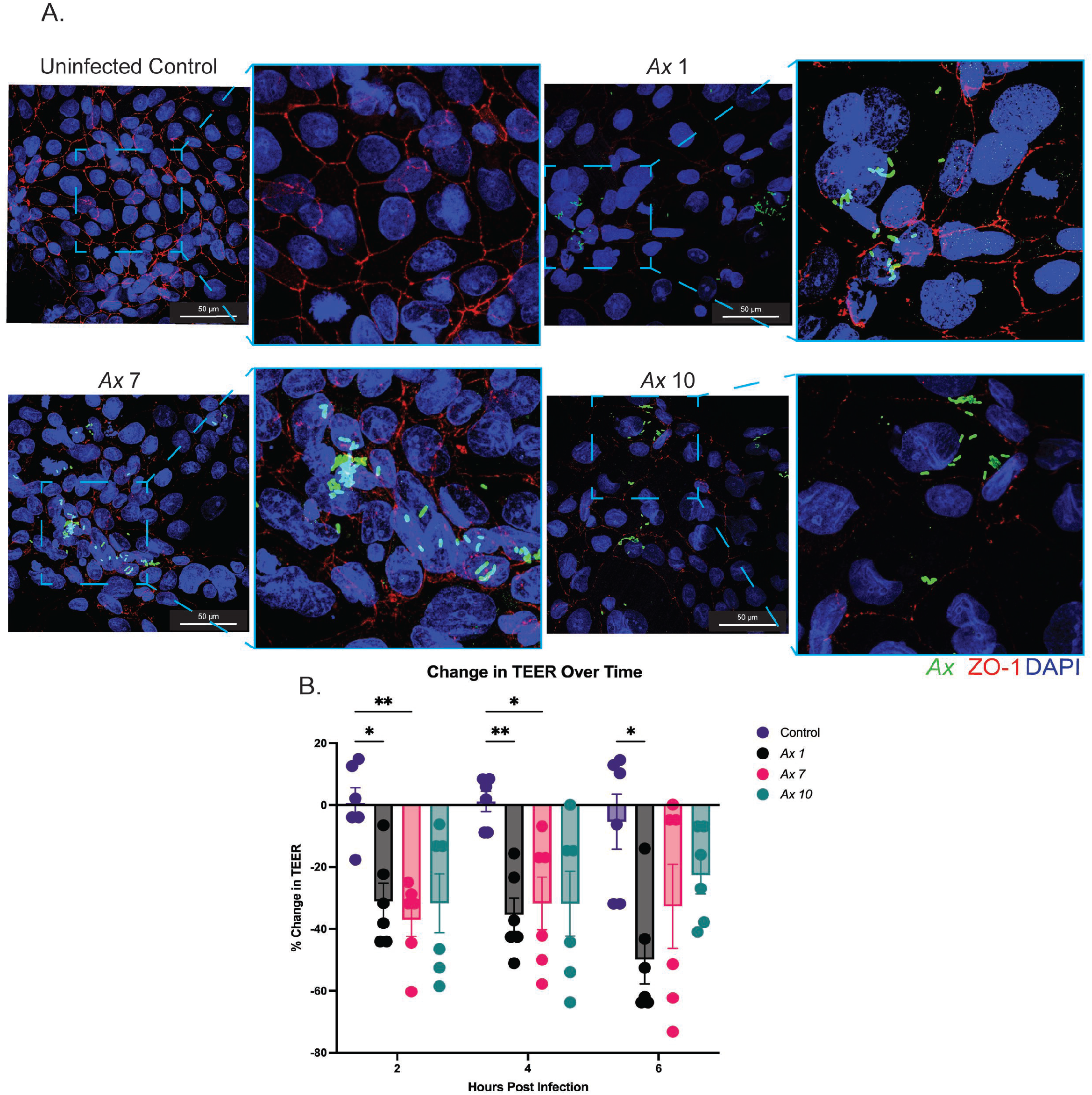
*Ax* strains compromise tight junction integrity of CFBEs grown at air-liquid interface. Submerged CFBE41o- (CFBE) cells were grown at air-liquid interface (ALI) were infected with MOI 10 of *Ax* effects on TEER and tight junction integrity were examined (n=6). A) 6 hour treated cells grown at ALI were stained to visualize tight junction integrity using an anti-ZO-1 antibody (red) and anti-*Ax* antibody (green). Cell nuclei were stained using DAPI (blue). Magnification= 60x; outlined sections were enlarged by 2x zoom. B) Transepithelial electrical resistance (TEER) was measured to examine tight junction integrity. Mean ± SEM. Two-way ANOVA with Tukey’s multiple comparisons test for post hoc analysis.* P<0.05, ** P<0.01, *** P<0.001, **** P<0.0001.

### *A. xylosoxidans* colonization and persistence in experimental mouse respiratory infections

After observing pathogenic phenotypes *in vitro*, we then wanted to examine the capacity for these isolates to colonize and persist in the murine lung. We intratracheally inoculated BALB/cJ mice with ∼10^7^ CFUs of each *Ax* strain. Mice were euthanized at 24, 48, and 72 hours post infection. We observed that each strain colonizes the murine lung, and we recovered at or near inoculum CFUs for each strain at the 24-hour time point (Fig. 3A). CFUs were slightly lower for each strain at later timepoints but were still well above our limit of detection (100 CFU/lung) (Fig. 3A). In addition, the amount of *Ax* adherent bacteria was not significantly different, indicating that each of these strains colonizes and persists similarly, despite differences in adherence to epithelial cells observed (Fig. 3A). Despite similarities in *Ax* bacterial load, there were differences in mouse virulence and mortality between these strains. *Ax* 7 and *Ax* 10 caused more mortality than *Ax* 1, with *Ax* 10 exhibiting a mortality rate of 80% at the 72-hour time point (compared to 0% for *Ax* 1 and 20% for *Ax* 7) (Fig. 3C). Despite differences in mortality, we did not observe significantly different weight change post infection between infection groups (Fig. 3B). Each infection group, however, did lose significantly more weight than the mock-infected controls (**** P <0.0001). After observing colonization and persistence in the murine lung, we examined how *Ax* was localizing in the lung tissue. We IF stained fixed lung sections using an anti-*Ax* antibody and DAPI to visualize lung cells. We observed *Ax* in the murine lung, where each strain of *Ax* appears to form aggregates (Fig. 3D).

**Figure 3:**
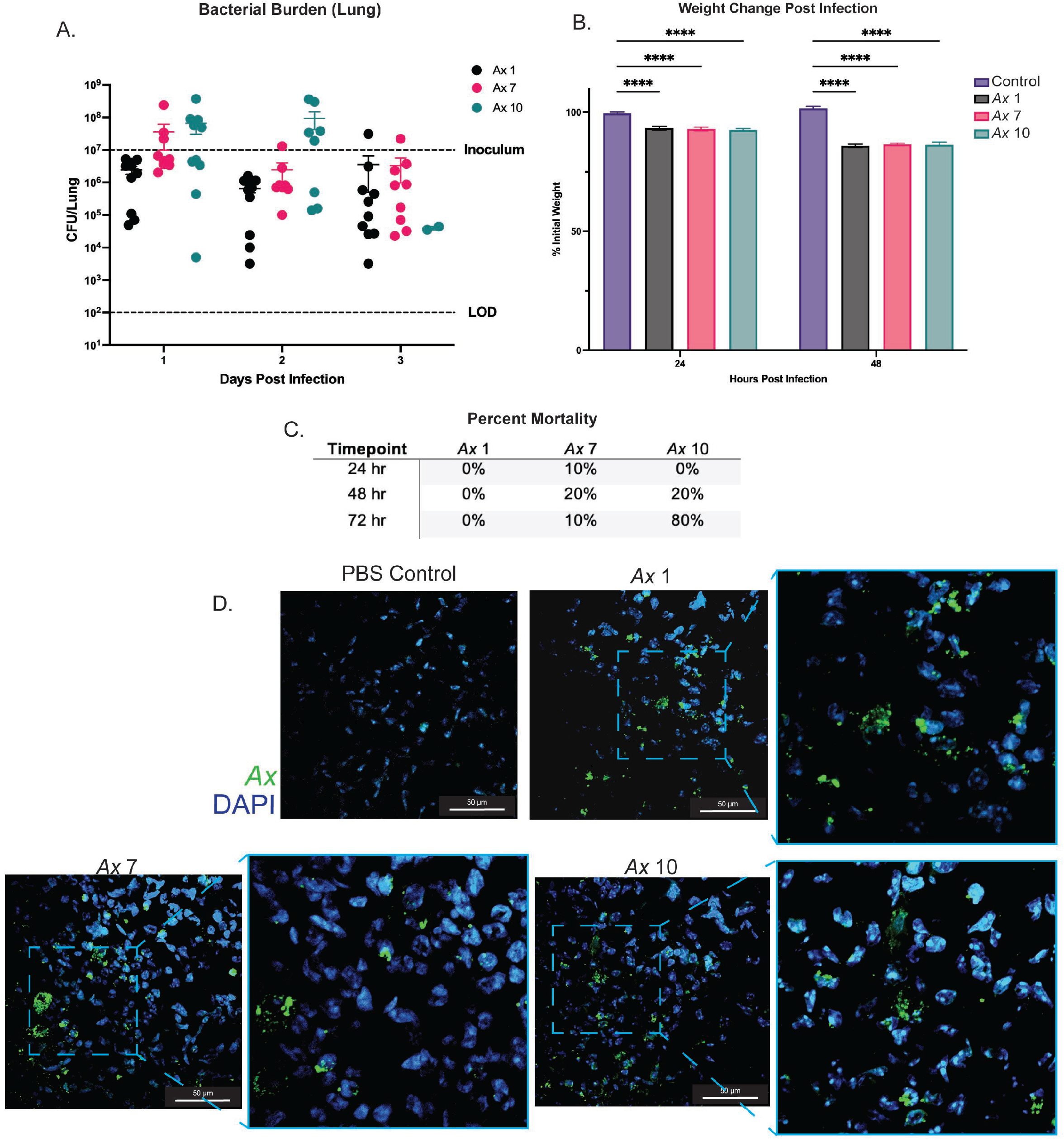
*Ax* strains colonize and persist in the murine lung up to 72 hours post infection. BALB/cJ mice were intratracheally infected with *Ax* 1, 7, and 10. A) *Ax* strains colonize and persist in the murine lung up 72 hours post infection. Mean ± SEM. Two-way ANOVA with Tukey’s multiple comparisons test for post hoc analysis. * P<0.05, ** P<0.01, *** P<0.001, **** P<0.0001. B) Percent mortality for each infection group for each *Ax* strain examined. C) Immunofluorescent (IF) staining of *Ax* in mouse lungs at the 48 hour time point. Sections were stained using an anti- *Ax* antibody (green) and DAPI (blue). Magnification=60x; outlined sections were enlarged with 2x zoom.

### Lung infection with *A. xylosoxidans* isolates causes lung damage and neutrophilia

After infecting mice with *Ax* isolates, we examined the immune response to infection with different *Ax* strains. We collected bronchoalveolar lavage fluid (BALF) from mice to perform differential cell counts on immune infiltrate and to quantify cytokines. Total cell counts from the BALF were similar between groups, except for *Ax* 7 (Fig. 4A). The BALF from *Ax* 7 infected animals had significantly more immune cell infiltration than each of the other groups as determined by the number of cells at 24 hours (**** P <0.0001) (Fig. 4A). At the 48-hour timepoint, however, the total cell counts from BALF for *Ax* 7 infected animals were only significantly higher than the mock-infected PBS controls (* P <0.05) (Fig. 4A). As expected from literature surrounding acute infection with Gram-negative pathogens, we observed that ∼80- 90% of the immune cell composition for *Ax* infected mice was composed of neutrophils (PMNs) (Fig 5A). Neutrophil composition of the BALF is not statistically different between each *Ax* strain but is statistically higher than mock infected controls (**** P <0.0001) (Fig. 4A). In addition, each infection group had comparable counts of macrophages in the BALF with the mock-infected groups having significantly higher percentage of macrophages present (****P <0.0001 at 24 hours post infection and **P <0.01 at 48 hours post infection), as expected due to the presence of resident lung macrophages (Fig. 4C).

**Figure 4:**
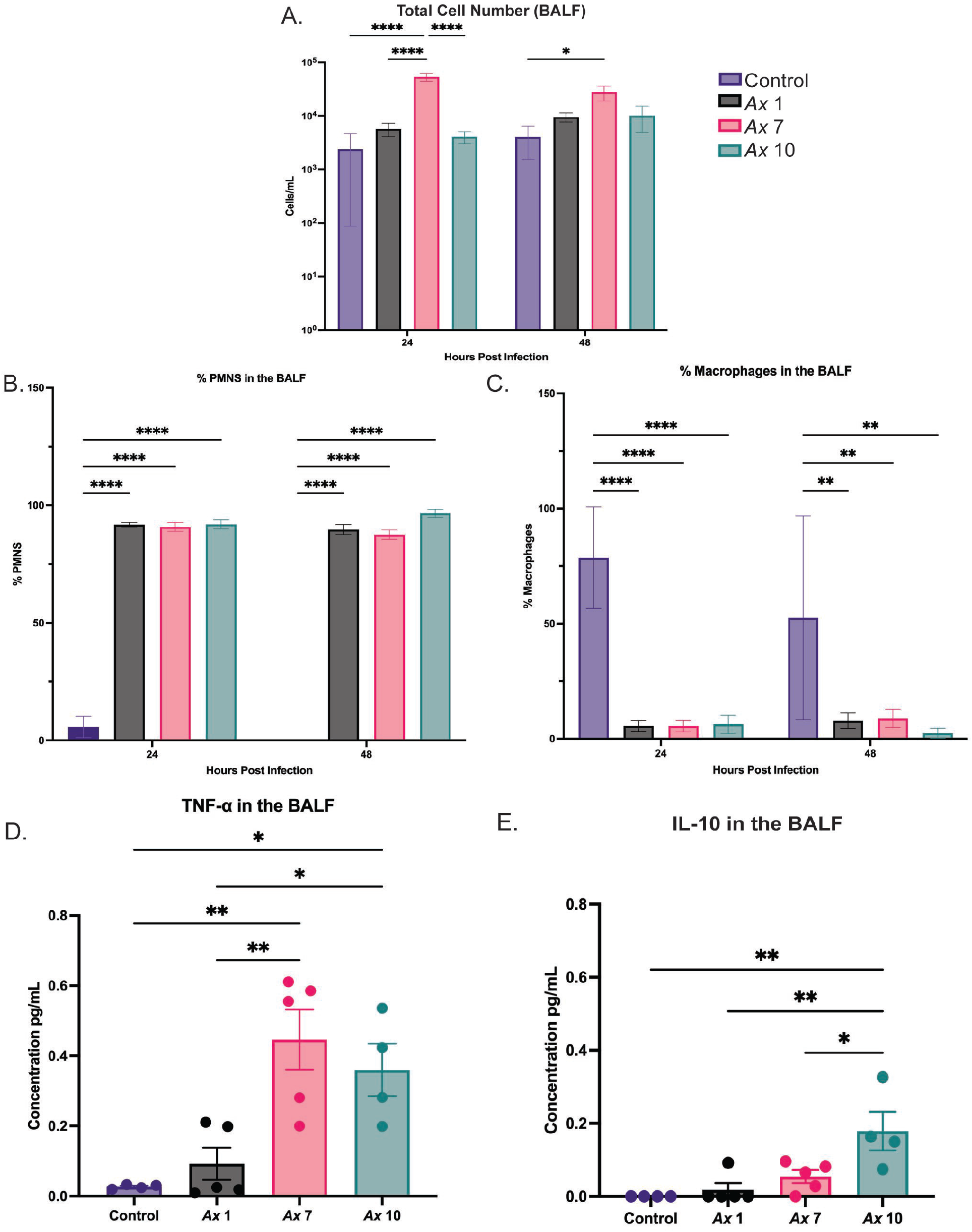
Immune response to infection with *Ax* strains. BALB/cJ mice were infected with *Ax* 1, 7, and 10 to assess the immune response post infection (n=4-5). A) Total leukocyte influx in the BALF 24 and 48 hours post infection. Mean ± SEM. Two-way ANOVA with Tukey’s multiple comparisons test for post hoc analysis. B) Percent of polymorphonuclear cells (PMNs) in the BALF of mice. Mean ± SEM. Two-way ANOVA with Tukey’s multiple comparisons test for post hoc analysis. C) Percent of macrophages in the BALF of mice. Mean ± SEM. Two-way ANOVA with Tukey’s multiple comparisons test for post hoc analysis. D) Quantification of TNF-α in the BALF at 48 hours post infection. Mean ± SEM. One-way ANOVA with Tukey’s multiple comparisons test for post hoc analysis E) Quantification of IL-10 in the BALF 48 hours post infection. Mean ± SEM. One-way ANOVA with Tukey’s multiple comparisons test for post hoc analysis. * P<0.05, ** P<0.01, *** P<0.001, **** P<0.0001.

**Figure 5:**
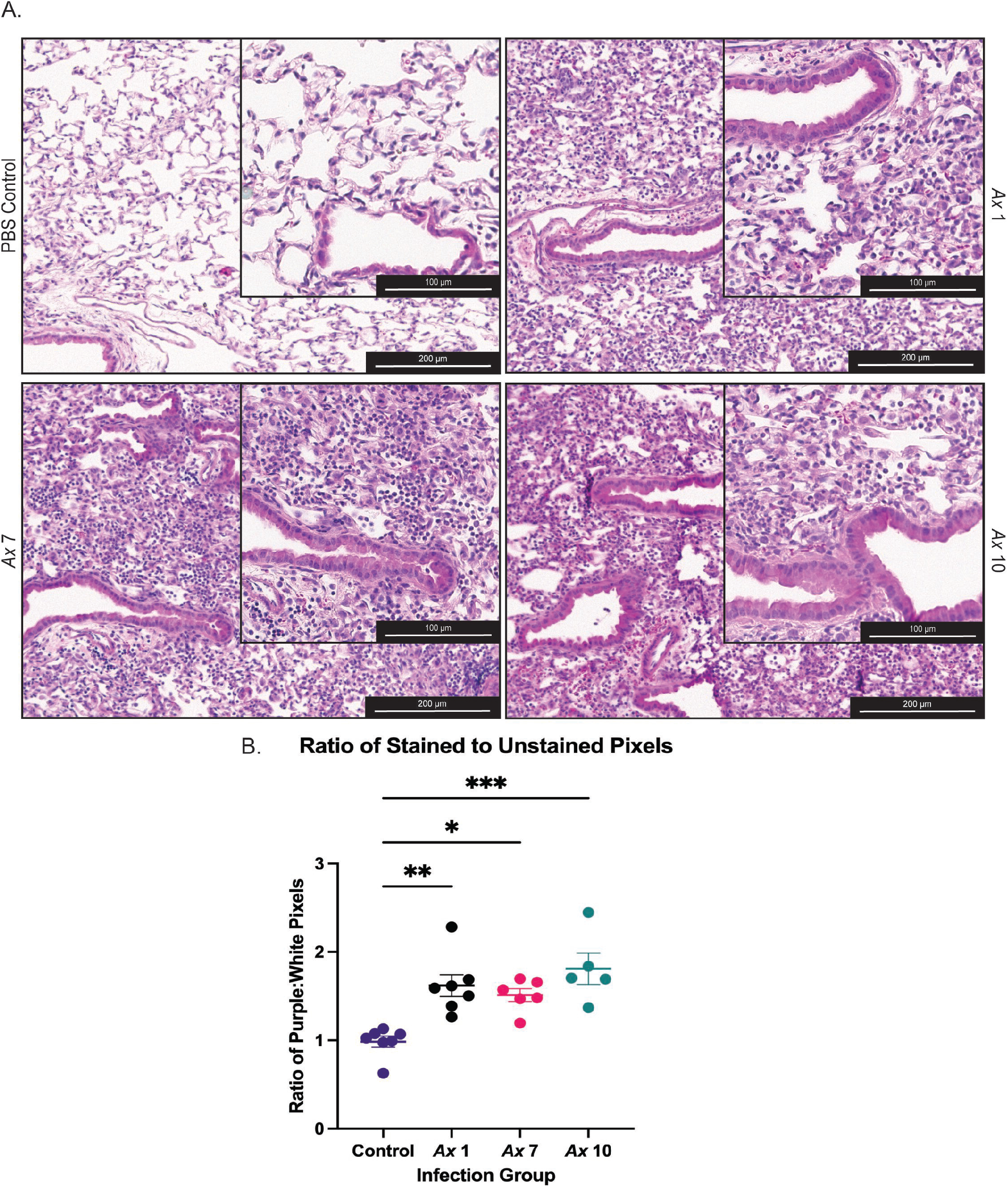
Histological changes in response to infection with *Ax* strains. Mice were infected with each strain of *Ax* and mock-infected with PBS. Animals were euthanized 48 hours post infection and the right lung was inflated and trimmed for hematoxylin & eosin (H&E) staining (n=5-8). A) Representative images of H&E stained *Ax* infected lungs 48 hours post infection. Magnification=10x and 20x (for the inlay). B) ImageJ was used to measure the pixel area stained (purple pixels) and unstained (white pixels). Points are an average the ratio of purple: white pixels for 3 images per animal. Mean ± SEM. One-way ANOVA with Tukey’s multiple comparisons test for post hoc analysis. * P<0.05, ** P<0.01, *** P<0.001, **** P<0.0001.

Following differential cell counting, we then examined whether there were differences in cytokine concentrations between mice infected with different strains of *Ax*. BALF from mice infected with *Ax* 1 did not have statistically higher concentrations of TNF-*α* than the control mock-infected mice (Fig. 4D). BALF from mice infected with *Ax* 7 and 10, however did have statistically different concentrations of TNF- *α* when compared to control and *Ax* 1 infected mice, but not when compared to each other. Furthermore, we observed significantly higher concentrations of IL-10 in *Ax* 10 infected animals than *Ax* 1 infected (** P <0.01), *Ax* 7 infected (* P<0.05), and control mice (** P<0.01) (Fig. 4E). *Ax* 1 and *Ax* 7 infected animals did not have significantly higher levels of IL-10 than the control mice (Fig. 4E).

In order to assess histopathologic changes, lung sections were stained with hematoxylin and eosin (H&E) and examined by light microscopy. Three representative images per animal were taken for quantification of inflammation and lung damage. The ratio of stained to unstained pixels was significantly higher for each infection group when compared to the controls (Fig. 5B), with the difference between *Ax* 10 and the control being the most statistically significant (***P<0.001). Additionally, the average ratio of stained to unstained pixels for each infection group were not statistically different from each other, indicating that the histopathological consequences for infection with *Ax* do not differ between these strains. An increased ratio of stained to unstained pixel area indicates that consolidation is occurring, in which the lung tissue is becoming denser due to inflammation and lung damage. This can be observed in the representative images (Fig. 5A), in which the tissue infected groups appear to be denser and more crowded. Additionally, other signs of lung damage appear such as increased neutrophilic influx (as confirmed by the increased neutrophils in the BALF), and airway thickening, which indicate increased inflammation and lung damage when compared to control mock-infected mice (Fig. 5A).

### *A. xylosoxidans* exhibits differential gene expression in artificial cystic fibrosis sputum media

After examining differential pathogenic phenotypes *in vitro* and *in vivo*, we wished to examine gene expression of *Ax* when grown in an artificial synthetic sputum media that mimics the CF lung environment (SCFM2) to identify genes that may be important for survival and pathogenesis in the CF lung. For this analysis, we chose to test the strain *Ax* 7, as it is closely related to MN001, a clinical isolate from another study, and it is intermediately pathogenic compared to *Ax* 1 and *Ax* 10 from our infection data (14). We grew *Ax* 7 in LB and SCFM2 to compare gene expression of *Ax* in these two media and identify genes that may be upregulated by *Ax* strains in the CF lung environment. RNA was extracted for sequencing from samples grown in these two media. Principle component analysis of these samples shows that those grown in LB cluster together, indicating that these samples have similar gene expression profiles that are also distinct from those samples grown in SCFM2 (Fig. 6A). Those grown in SCFM2 also appear to cluster together, except for one sample, which still does not cluster with those grown in LB (Fig. 6A). Differential expression analysis showed that there were 435 significantly differentially regulated genes between *Ax* 7 grown in LB and SCFM2 medium. Additionally, a heatmap that showing the top 35 most differentially expressed genes confirms that gene expression profiles are similar between samples in the same treatment group (Fig. 6B). Pathway analysis via ClusterProfiler was used to identify enriched KEGG pathways between these two groups. Compared to gene expression in LB, there were 12 statistically different enriched KEGG pathways among *Ax* grown in SCFM2, with 6 of these being upregulated and 6 of these being downregulated (Fig 6D). Of the upregulated KEGG pathways, there was quorum sensing, ABC transporters, fatty acid metabolism, degradation, and biosynthesis, and nitrogen metabolism (Fig 6D). The KEGG pathways that were significantly downregulated in the SCFM2 group were flagellar assembly, aminoacyl-tRNA biosynthesis, purine metabolism, biosynthesis of cofactors, nucleotide metabolism, and porphyrin metabolism (Fig 6D).

**Figure 6:**
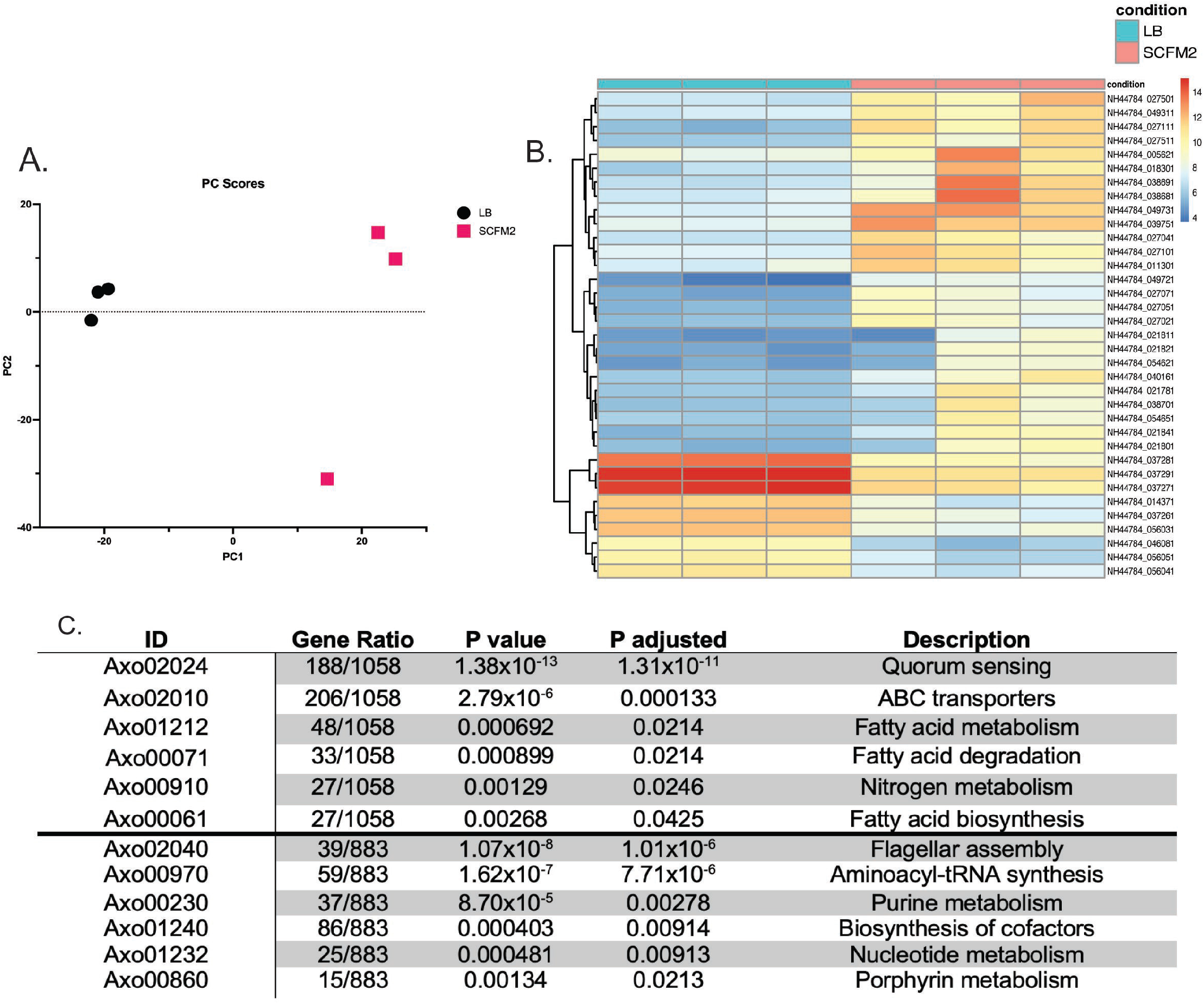
Differentially expressed genes from *Ax* grown in LB and SCFM2. *Ax* 7 was grown in lysogeny broth (LB) and a synthetic CF sputum media (SCFM2). RNA was extracted and sequenced for differential expression analysis. A) Principal component analysis of *Ax* 7 grown in LB and SCFM2. B) Heatmap of the top 35 variably expressed genes between *Ax* 7 grown in LB and SCFM2. C) The 6 most highly significantly upregulated (top) and downregulated (bottom) KEGG pathways among samples grown in SCFM2 compared to those grown in LB.

## DISCUSSION

Respiratory infections are a major cause of morbidity and mortality for pwCF (18). There remains much to be learned regarding the clinical significance of specific microbes in pwCF and this is particularly unclear for emerging pathogens such as *A. xylosoxidans*. *A. xylosoxidans*, has been increasingly observed in sputa and other clinical samples from pwCF. In this study, we aimed to examine virulence determinants present in clinical *Ax* strains and to begin to examine and characterize the consequences of pulmonary infection with *Ax*. Herein, we show that *Ax* strains isolated from pwCF possess and express virulence factors to a different extent but colonize and persist in a murine model of infection similarly. In addition, we observed that *Ax* strains produce different immune responses during murine pulmonary infection despite recovering similar CFUs amongst infection groups. Furthermore, we examined differences in gene expression profiles between an *Ax* strain grown in LB compared to growth in media mimicking the CF lung environment, indicating that *Ax* upregulates certain metabolic and virulence pathways in the CF lung.

Previous genomic sequence profiling of *Ax* strains have demonstrated orthologs for genes associated with capsular polysaccharide, cytotoxicity, motility, and biofilm formation/maturation (9, 11). We began with an analysis of the genomic content of *Ax* strains and an examination of the virulence related genes that these strains possess. We saw that our *Ax* strains harbor genes associated with several virulence factors such as flagellum and chemotaxis, type IV secretion, adherence, and antibiotic resistance (data not shown). Phenotypic analysis showed that despite possessing similar virulence genes, *Ax* isolates exhibit differing phenotypes *in vitro*. In this study, we observed that all the isolates examined possess flagellar genes, but only *Ax* 10 exhibits swimming motility, which is mediated by the presence of flagella (Fig. S1B). This discrepancy between genotype and phenotype may be explained by a phenomenon known as hypermutation. This is well studied in *Pseudomonas aeruginosa* (*Pa*) and is understood to contribute to differential gene expression in strains that are chronic colonizers (19–21). Hypermutation occurs due to mutation in mismatch repair systems that allow for bacteria to rapidly mutate genes and respond to selective pressure (20). Although it is not well-studied in *Ax*, hypermutation has been observed in this species and may be a factor that contributes to the discrepancies between genotype and phenotype that we have observed here (22, 23).

To test colonization, persistence, and virulence of *Ax* in the context of pulmonary infection, we carried out a series of murine infection studies. Previous infection studies have been done in mice, but they were via intraperitoneal infection in immunocompromised mice (24). With our murine pulmonary infection model, we observed that *Ax* strains colonize and persist in the lungs for up to 72 hours post infection, with there being no significant difference in CFUs at each timepoint for each strain (Fig. 3A). Previous work from our lab shows that a similar late-stage Gram-negative CF pathogen, *Stenotrophomonas maltophilia*, does not colonize and persist as well as our *Ax* isolates in a similar single infection model (25). We also began to characterize the immune response to *Ax* infection and observed that there are immunological consequences including higher levels of inflammatory cytokines, neutrophil influx, lung damage, and mortality (Fig. 6), which may contribute to disease progression and lung damage in pwCF.

Prior work has shown that TNF-*α* levels are increased in the sputum of *Ax* infected pwCF (26). We quantified TNF-*α* and an anti-inflammatory cytokine, IL-10, in BALF from mice infected with *Ax* and uninfected controls. Levels of IL-10 and TNF-a showed variable increases following infection, with most of the Ax infected mice yielding significantly more cytokine than uninfected controls (Fig. 4). Increased cytokine levels in the lung are well understood in CF and cause increased neutrophil infiltration, which leads to lung damage (2, 3).These data indicate that *Ax* strains can produce differing immune responses and can potentially have different effects on CF disease progression and lung damage, depending on the extent of the immune response induced by each strain. In addition, despite recovering comparable *Ax* bacterial load, our animal data showed different levels of cytokine production and mortality between *Ax* strains, further exemplifying differences in pathogenicity between *Ax* strains and the lung damage induced by these strains.

Furthermore, we examined growth and gene expression amongst a CF isolate of *Ax* in the presence of SCFM2, a medium that was designed to mimic the CF lung environment (27, 28). We observed that compared to growth in LB, *Ax* 7 upregulates metabolism genes in SCFM2. Interestingly, we also observed that *Ax* 7 upregulates genes associated with quorum signaling. These results indicate genes that may be necessary for growth in the CF lung environment, but more importantly, they indicate that *Ax* may upregulate quorum sensing genes in this environment, which are essential for inducing many pathogenic processes (29, 30). Quorum signals allow for bacteria to communicate with each other (either as a species or between species) and induce expression of specific genes when bacterial density and the concentration of signal molecules reaches a certain point (30). Many important pathogenic genes have been known to be under the control of such pathways, notably biofilm formation and toxin secretion (29, 30). This finding indicates that quorum signaling may not only be important for *Ax* pathogenesis, but specifically for *Ax* colonization and persistence in the CF lung environment. Additionally, an increase in quorum sensing gene expression may also indicate that *Ax* is primed for communication with other bacterial species in the CF lung, as these infections are polymicrobial and complex (31).

Downregulation of flagellar assembly genes indicate that flagellum-mediated motility may not be an important virulence determinant for *Ax* in the CF lung. This combined with the lack of flagellar mediated motility in this strain further implies that this may not the most important virulence determinant for *Ax*, especially in the context of CF. Other enriched genes are mainly associated with metabolism, which gives us insight to which metabolites *Ax* can take advantage of in the CF lung environment. Additionally, this may be informative with regards to polymicrobial interactions that take place in CF, as the CF lung is colonized by more than one microbial species at once (32, 33). A factor that determines the extent of polymicrobial interactions, especially in the host where nutrients are often limited, is nutrient acquisition (34–36). The fact that *Ax* can metabolize or may preferentially metabolize certain compounds could give it a competitive edge against other microbial species in the CF lung.

Carriage of *Ax* in pwCF has been established and steadily increasing since the 1980’s (8–10). Clinical studies have shown that *Ax* is an opportunistic pathogen that causes disease in a variety of contexts, but there is a need for examination of its pathogenicity in pulmonary infection (8, 37). Recent studies have examined genes important for biofilm formation and cytotoxicity to macrophages, but these studies are largely *in vitro* or in non-pulmonary cell lines (14–16). Our work establishes a murine pulmonary infection model for *Ax* and shows that strains can colonize and persist in the lung for up to 72 hours post infection. In addition, we saw increased levels of cytokines, neutrophil infiltration, lung damage, and mortality in our infection model. We also identified genes that were differentially expressed in *Ax* when grown in a medium that mimics the CF lung environment, suggesting that *Ax* strains may increase expression of some pathogenic genes in the CF lung. Overall, we found that *Ax* has pathogenic properties *in vivo*, and that *Ax* may have significant effects on CF disease progression. Ongoing work is occurring in our laboratory to identify specific genes and their role in pathogenesis in a mammalian host.

## MATERIALS AND METHODS

### Whole genome sequencing and analysis

Bacterial strains were grown overnight in broth cultures (37° C, shaking at 200 RPM) and had genomic DNA extracted using a DNeasy Blood and Tissue kit (Qiagen). Genomic DNA was sequenced using the Illumina MiSeq at the Heflin Genomics Core at UAB. Reads were trimmed and assembled into contigs via PATRIC.

### Bacterial strains and growth conditions

*Ax* 1, 7, and 10 are sputum-derived isolates from pwCF provided by Timothy Starner (University of Iowa Children’s Hospital). Strains were cultured by streaking for colony isolation onto lysogeny broth (LB) with agar (Difco) and incubating overnight at 37° C. Single colonies were used to inoculate overnight cultures of LB (37° C, shaking at 200 RPM). For experiments, overnight cultures were diluted to the appropriate optical density at 600 nm (OD_600_) for ∼10^8^ CFUs and diluted further if needed.

### Crystal violet biofilm assay

Overnight cultures were diluted and seeded at ∼10^8^ CFUs in a 96 well plate. Biofilms were incubated at 37°C without shaking. At each time point (4, 6, 8, and 24 hours), cultures were washed and either stained with crystal violet or plated for viable colony counting. Crystal violet staining was performed based on a previously published protocol (38).

### Motility assays

Swimming motility assays were adapted from a previously published protocol for *Pseudomonas aeruginosa* (39). We made low percentage (0.3% agar) LB plates. Plates were poured on the day of the assay and sat for 4 hours before inoculation. Overnight cultures were inoculated halfway into the agar with a toothpick. Plates were incubated upright at 37°C and imaged 24 and 48 hours post inoculation. Twitching assays were done 1.5% agar LB plates based off a previously published protocol (40). Plates were poured on the day of the assay and sat for three hours before inoculation. Holes were stabbed into the bottom of the agar (touching the plate) with toothpicks. Then, we inoculated plates with toothpicks that were dipped in an overnight culture of each strain (the bottom of the plate). Plates were incubated upright at 37°C. After 24 and 48 hours post inoculation, the agar was removed, and the plates were stained with 0.1% crystal violet to visualize the area covered by bacteria. We took images and measured the diameter of the area traveled in ImageJ.

### Mammalian epithelial cell culture

Immortalized human cystic fibrosis bronchial epithelial cells (ΔF508/ΔF508 CFBE41o-) were maintained in minimal essential medium (MEM) supplemented with 10% fetal bovine serum (FBS) (41). For submerged cell cultures, cells were seeded at ∼10^5^ cells per well. For air-liquid interface (ALI) cell experiments, filter membranes were seeded with ∼5 x 10^5^ cells per well, and liquid was kept on the apical membrane surface for one week and removed to leave cells at an air-liquid interface (ALI). After one week at ALI, cells were infected as indicated in the relevant figure legends. For all cell experiments, cells were infected with a multiplicity of infection (MOI) of 10 bacterial cells per mammalian cell. After infection, cells were washed twice with sterile PBS to remove non-adherent bacteria and either plated or fixed for staining.

### Immunofluorescent staining

CFBE cells were grown at ALI and infected with an MOI of 10 of each bacterial strain for 6 hours, after which the cells were fixed in ice cold methanol-acetone (1:1) and immuno- fluorescently stained using a previously published protocol (42, 43). Cryosectioned lung tissue sections (5 μm/section) from infected mice were stained using polyclonal anti-*Ax* rabbit sera and appropriate fluorescent conjugate secondary antibody (Invitrogen). Tight junctions were stained via a polyclonal goat antibody against ZO-1 and relevant secondary antibody (Invitrogen). DAPI was used to stain host cell nuclei. Stained cells were imaged on a Nikon-A1R HD25 confocal laser microscope.

### Confocal microscopy

Confocal laser scanning microscopy (CLSM) was performed using a Nikon-A1R HD25 microscope (Nikon, Tokyo, Japan). Images were acquired and processed using the NIS-elements 5.0 software.

### Mouse model of respiratory infection

BALB/cJ mice (8 to 10 weeks old) were obtained from Jackson Laboratories (Bar Harbor, ME). Mice were anesthetized with isoflurane and intratracheally inoculated with *Ax* (∼10^7^ CFU in 100 µL PBS). For viable plate counting, the left lung of each mouse was homogenized in 500 µL of sterile PBS at 30 Hz/s. The resulting lung homogenate was serially diluted in PBS and plated on LB agar to obtain viable colony counts. The right lung was inflated with 10% buffered formalin and stored at 4µC until processing for sectioning and histopathological analysis. Mock-infected control mice were intratracheally inoculated with 100 μL of sterile PBS and processed in the same manner.

### Bronchoalveolar lavage fluid (BALF) collection

Bronchoalveolar lavage was performed by flushing lungs with 6 mL of sterile PBS in 1 mL increments. Collected BALF was stored on ice until processing. The first mL isolated was centrifuged and resuspended in fresh PBS to be mounted on a slide via cytospin at 500 rpm for 5 minutes. The remaining 5 mL of BALF was stored at −20°C until processing.

### Cytokine analyses and differential cell counting

Cytokine analyses were performed on the 5 mL of collected BALF via DuoSet ELISA kits (R&D Systems, Minneapolis, MN). For total and differential cell counts, cells mounted via cytospin were stained with a Kwik-Diff differential cell stain (Fisher Scientific, Waltham, MA). Three representative fields per view per sample were counted to determine the composition of immune cells.

### Histological analysis

Right lungs were inflated in 10% buffered formalin (Fisher Scientific, Waltham, MA) and stored at 4°C until processing. Each lobe of the right lung was trimmed, and sections were sent to the UAB Comparative Pathology Laboratory for paraffin embedding, sectioning, and hematoxylin and eosin (H&E) staining. For quantification, three images per mouse were taken and analyzed using ImageJ. Stained to unstained pixel area ratios were averaged from the three photos per animal.

### Gene expression in SCFM2

Gene induction experiments were performed using a synthetic CF sputum medium (SCFM2), prepared as described by a previous publication (27). LB was used as a rich medium control for this experiment. 5 mLs of the respective growth medium were inoculated with ∼10^8^ CFU of *Ax* and due to difficulty in determining growth via optical density in SCFM2, cultures were plated for viable colony counting to examine growth over time. Samples were grown to ∼10^10^-10^11^ CFU/mL, as confirmed by plating. To prepare samples for RNA extraction, RNA-later in equal volume to the growth media was added to each sample. Samples were frozen at −80°C until processing.

### RNA extraction and sequencing

Samples were defrosted and spun down to remove residual media and RNA-later. Bacterial pellets were suspended in Trizol reagent (Invitrogen) and lysed by bead beating (0.1 mm silica). RNA extraction was performed using a standard protocol (44). Extracted RNA was then DNase treated (DNase I, NEB) and processed through another Trizol extraction to purify the example of enzyme. DNase treated samples were then sent to SeqCenter (Pittsburgh, PA) for library preparation and sequencing on the Illumina NextSeq2000 platform.

### Trimming, alignment, and differential expression analysis

Reads were trimmed with Trim Galore! (v. 0.4.4) and Cutadapt (v. 1.9.1) and evaluated for quality using FastQC. Reads were aligned to the published *A. xylosoxidans* strain NH44784- 1996 genome using STAR aligner (v. 2.7.3a). Read counts were obtained via Subread FeatureCounts and differential expression analysis was performed using DESeq2. Pathway analysis was performed using clusterProfiler (v. 4.4.1) with Gene Ontology and KEGG pathway databases.

### Statistical analyses

All graphs represent sample means ± the standard error of the mean (SEM) unless otherwise noted. All experiments were repeated at least two times and data from independent experiments were combined for analysis. All statistical tests were performed using GraphPad Prism 8 (San Diego, CA).

## ACKNOWLEDGEMENTS

The authors gratefully acknowledge helpful input and discussions with colleagues in the UAB Center for Cystic Fibrosis Research. This work was supported by grants from the Cystic Fibrosis Foundation (CFFSWORDS1810, CFF SWORDS20G0) awarded to W.E.S. as well as the CF Research and Translation Core Center grant (P30DK072482) and the UAB Research and Development Program (ROWE19RO). C.E.B. is supported by the American Indian Science and Engineering Society (AISES) Lighting the Pathway Program (LTP). We also thank Timothy Starner (University of Iowa Children’s Hospital) for bacterial strains used in this study.

## SUPPLEMENTAL FIGURE LEGENDS

**Supplemental Figure 1:** *In vitro* assays were performed to examine *Ax* phenotypes that have previously been described in other *Ax* strains, such as biofilm formation, motility, and antibiotic resistance. A) Biofilm formation on a static abiotic (plastic) surface exhibited by *Ax* strains as indicated by crystal violet (CV) staining. Mean ± SEM. Two-way ANOVA with Tukey’s multiple comparisons test for post hoc analysis. B) Swimming motility of isolates was examined by inoculation into low percentage agar plates. Distance was measured as the diameter of the area traveled. C) Twitching motility of isolates was examined by inoculation on the bottom of low percentage agar plates. Distance was measured as the average diameter of area travelled. Mean ± SEM. One-way ANOVA with Tukey’s multiple comparisons test for post hoc analysis. * P<0.05, ** P<0.01, *** P<0.001, **** P<0.0001.

